# Exploration and analysis of R-loop mapping data with *RLBase*

**DOI:** 10.1101/2021.11.01.466854

**Authors:** H. E. Miller, D. Montemayor, J. Li, S. Levy, R. Pawar, S. Hartono, K. Sharma, B. Frost, F. Chedin, A. J. R. Bishop

## Abstract

R-loops are three-stranded nucleic acid structures formed from the hybridization of RNA and DNA during nascent transcription. In 2012, *Ginno et al.* introduced the first R-loop mapping method, DNA:RNA immunoprecipitation (DRIP) sequencing. Since that time, dozens of studies have implemented R-loop mapping and new high-resolution techniques have been developed. The resulting datasets have tremendous potential to reveal the causes and consequences of R-loops genome-wide. However, poor quality and variability between mapping approaches pose serious barriers to the meta-analysis of these data. In our recent work, we reprocessed 693 R-loop mapping samples, devising new quality methods, defining a set of high-confidence mapping samples, and then deriving R-loop regions, consensus sites of R-loop formation. This analysis yielded the largest R-loop data resource to date along with novel computational approaches for R-loop mapping analysis. Now, we introduce *RLBase*, an innovative web server which builds upon those data and software by providing users with the capability to (1) explore hundreds of public R-loop mapping datasets, (2) explore consensus R-loop regions, (3) analyze user-supplied datasets to generate an HTML quality report, and (4) download all the processed data for the 693 samples we previously reprocessed and standardized. In addition to *RLBase*, we also describe the other software which, along with *RLBase*, provides a computational framework for R-loop bioinformatics. *RLBase*, and the rest of these software (termed “RLSuite”), are provided freely under an MIT license and made publicly available: https://gccri.bishop-lab.uthscsa.edu/rlsuite/. RLBase is directly accessible via the following URL: https://gccri.bishop-lab.uthscsa.edu/rlbase/.

## Introduction

R-loops are three stranded nucleic acid structures comprising an RNA:DNA hybrid and displaced single-stranded DNA (1). They occur as a byproduct of transcription at some genes, and can be promoted by a variety of factors, including high G/C skew (2) and negative DNA superhelicity (3). Previous studies have implicated R-loops in a variety of pathological consequences, including their potential to promote replication stress (4), a phenomena observed in hypertranscriptional cancers such as Ewing sarcoma (5–7). Recent evidence has indicated that R-loops may also play important physiological roles in gene expression (1), DNA repair (8), and ribosome biogenesis (9). However, the dynamics of R-loops, the mechanisms of their regulation, and the delineation between pathological and physiological R-loops remain largely unclear (1, 10).

R-loop mapping studies apply high-throughput sequencing (HTS) to examine the causes and consequences of R-loops under differing biological conditions. However, each study is limited by the narrow definition of the biological system which it examines and by the limitations of the techniques used in R-loop mapping (11). The mining of public datasets can reveal insights into the fundamental biology of a molecular system, as was previously demonstrated in the transcriptomic (12) and epigenomics (13) fields. Our 2020 study of R-loops in the context of chromatin architecture was, to our knowledge, the first to employ data mining to assess fundamental R-loop biology (14). While our profiling of 108 DRIP-sequencing samples provided insights into the locations and dynamics of R-loops, it also revealed inconsistencies in data quality between published studies (14). These findings were reinforced recently (15), suggesting that data quality inconsistencies are a barrier in the R-loop field, and raising questions regarding the validity of findings derived from R-loop mapping studies.

To address these issues, we recently undertook the reprocessing and standardization of 693 R-loop mapping datasets, the largest R-loop mining project to date (11). From these data, we developed a new R-loop mapping quality control methodology which shows excellent accuracy and validity (11). With these approaches, we successfully defined a high-confidence set of R-loop mapping samples, and we performed a meta-analysis from which we defined consensus “R-loop regions” (RL regions) (11). As a result of this work, we also developed a suite of software packages for the analysis of R-loop data, including *RLPipes*, a CLI tool for upstream data analysis (Bioconda), *RLHub*, an R package for accessing the processed data from (Bioconductor), and *RLSeq*, an R package for downstream analysis of R-loop mapping data (Bioconductor). With these software and data resources, generated from our previous work, we created *RLBase*, a user-friendly web server which provides the capability to (1) explore hundreds of R-loop mapping datasets, (2) explore R-loop regions, (3) analyze R-loop mapping datasets in the browser, and (4) download processed and standardized R-loop mapping data.

Though another R-loop web server was previously developed, *R-loopDB*, it is designed for the exploration of R-loop forming sequences (RLFS), so it contains few re-processed datasets and lacks the ability to analyze R-loop mapping datasets (16).

Taken together, *RLBase* and the other software and data sources described in the present work, termed “RLSuite”, represent a dramatic advancement in R-loop bioinformatics and a valuable resource for the R-loop field. “RLSuite” and *RLBase* can be accessed publicly at the following URL: https://gccri.bishop-lab.uthscsa.edu/rlsuite/.

## Methods

### Preliminary data curation

Prior to data curation, it was necessary to obtain basic genome information and genomic features. This included: (1) UCSC genome information, (2) R-loop forming sequences (RLFS) predictions, and (3) genomic annotations tailored for R-loop analysis. The following subsections detail the processing steps involved in obtaining these data.

#### Genome metadata

Prior to running the data generation pipeline, it was necessary to curate a list of all available genomes in the UCSC data repository. For each genome in UCSC, the following information was obtained via the *makeAvailableGenomes.R* script in the *RLBase-data* GitHub repository (See *Code Availability*):

1. UCSC organism assembly ID
2. Taxonomy ID
3. Scientific Name
4. Year that the genome assembly was introduced
5. Genome length
6. Gene annotation availability (TRUE/FALSE)
7. Effective genome size at various read lengths (calculated via the *Khmer* python package).

The resulting data was then packaged for use with *RLPipes* and *RLSeq*.

#### R-loop forming sequences (RLFS)

R-loop forming sequences were developed via the methods we described previously (11).

#### Genomic annotation database

Finally, we developed a custom genomic annotation database for use with *RLSeq* that contained annotations relevant to R-loop biology. This was accomplished via a custom R script, *getGenomicFeatures.R*, in the *RLBase-data* GitHub repository (see *Code Availability*). For the full list of data sources and descriptions, see **Table S1**.

### RLBase data (upstream)

The processed data stored in *RLBase* and used by all other *RLSuite* software was generated using a long-running computational pipeline available in its entirety in the *RLBase-data* GitHub repository (see *Code Availability*). The steps for upstream data processing were described in detail in our recent work (11). In addition to those details previously described, the present work will also describe the process of analyzing RNA-Sequencing datasets.

#### RNA sequencing datasets

In cases where R-loop mapping studies also provided matched RNA sequencing (RNA-Seq) datasets, they were downloaded and quantified to assess gene expression with *RLPipes* via the following procedure: raw reads were downloaded and pre-processed using the same procedure described in our recent work (11) up until the genomic alignment step and then reads were quantified using the *salmon* pseudo aligner (17) to generate read counts.

### RLBase data (downstream)

Following generation of processed data files (peaks, coverage, expression quantification, and quality statistics), downstream data processing was initiated to generate the final *RLBase* data. This involved (**1**) R-loop forming sequences (RLFS) analysis, (**2**) quality model building, (**3**) sample classification, (**4**) R-loop consensus analysis, (**5**) peak annotation and enrichment testing, (**6**) Expression matrix generation, (**7**) R-loop region annotation, (**8**) sample-level correlation analysis, (**9**) R-loop region abundance matrix generation, (**10**) calculating R-loop/expression correlation, (**11**) updating RLHub, (**12**) updating the *RLBase* genome browser trackhub, (**13**) RLSeq analysis of every sample, (**14**) upload of all data to the RLBase AWS S3 bucket.

Notably, many of these steps (**1-5**, **8**) are described in detail in our previous work (11). Therefore, the present work will describe only those **(6-7**, **9-14**) which have not been described elsewhere.

#### Gene expression compilation

As part of the RLBase processing pipeline, RNA-Seq data from R-loop mapping studies was also analyzed. These data were quantified and saved to the disk (see *RLBase data (upstream)*). Then, the *buildExpression.R* script from the *RLBase-data* GitHub repository (see *Code Availability*) was executed to convert these quantification files to a *SummarizedExperiment* object, from the *SummarizedExperiment* R package (18). The procedure implemented in *buildExpression.R* followed these steps: (1) all quantification files were imported using the *tximport* function from the *tximport* R package (19) and summarized to the gene level. (2) The transcripts per million (provided in each file) was log2 transformed. (3) The variance-stabilizing transform (VST) of the data was calculated via the *vst* function from the *DESeq2* R package (20). The count matrix, log2TPM matrix, and VST matrix were then combined using the *SummarizedExperiment* function (18) and saved to disk.

#### R-loop Region annotation

R-loop regions (RL regions) were annotated via the *rlregionsToFeatures.R* script in the *RLBase-data* repository (see *Code availability*). For catalytically dead RNase H1 (dRNH), S9.6, and combined RL regions, the peaks were annotated with the hg38 genomic annotations previously described (see *Genomic features*) using the *bed_intersect* function from the *valr* R package (21).

#### R-loop region abundance calculation

R-loop region (RL region) abundance was calculated within each human sample in *RLBase* using the *rlregionCountMat.R* script from the *RLBase-data* GitHub repository (see *Code availability*). *RLBase* sample alignment files (“BAM” format) were processed with *featureCounts* from the *Rsubread* R package (22) to quantify the read counts from each within RL regions. Then log2 RL regions per million (log2RLRPM) was calculated using the same procedure for log2 transcript per million (log2TPM) calculation during gene expression analysis. Finally, the variance stabilizing transform (VST) was used to calculate normalized counts via the *vst* function from *DESeq2* (20). The three resulting matrices, raw counts, log2RLRPM, and VST, were combined into a *SummarizedExperiment* object using the *SummarizedExperiment* R package (18) and saved to disk.

#### R-loop region abundance correlation with gene expression

Next, the *rlExpCorr.R* R script from the *RLBase-data* GitHub repository (see *Code availability*) was used to calculate the correlation of R-loop region (RL region) abundance. The procedure used is describe below.

##### Matching R-loop samples and expression samples

Some R-loop mapping studies also had matched RNA-Seq data. For each R-loop mapping sample, the “study”, “tissue”, “genotype”, and “other” columns from the curated metadata were compared to the same column in the list of expression samples. If the values in all four columns were a match with at least one expression sample, then those four columns would be assigned as the “exp_matchCond” (expression match condition). If only three were available, then they would become the “exp_matchCond”. To see the order in which columns were checked for possible matches, view the *buildExpression.R* script in the RLBase-data GitHub repo (see *Code availability*).

##### Summarizing results within match conditions

Once match conditions were found, the R-loop abundance and gene expression data were summarized within them. Briefly, the R-loop abundance (log2RLRPM) was averaged within any R-loop mapping samples sharing the same “exp_matchCond” to create one summarized log2RLRPM value within each exp_matchCond – RL Region pairing. For the expression samples, genes were mapped to RL regions and the log2TPM was summed within them to create an RL region expression matrix. Then, the log2TPM within RL regions was averaged within any expression samples sharing an “exp_matchCond” to yield one summarized log2TPM value within each exp_matchCond – RL Region pairing. Then, the Spearman correlation was calculated between log2RLRPM and log2TPM across exp_matchCond within each RL region to yield a correlation estimate (*Rho*) and p value within each RL region. P value adjustment was performed using the Benjamini Hochberg procedure.

#### Finalizing RLHub data

*RLHub* is the *ExperimentHub* (23) package used to conveniently access the process data generated as part of the *RLBase-data* workflow (see *RLHub*). In this step of the workflow, processed data were packaged for *RLHub* using the *prepRLHub.R* script in the *RLBase-data* GitHub repository (see *Code availability*). The script was run, and the resulting data was uploaded to the *RLHub* directory within the *RLBase* AWS S3 bucket.

#### Building the RLBase genome browser session

The UCSC genome browser (24) provides a user-friendly way for exploring genomic data in a web browser interface. A TrackHub is a collection of genomic data that can be visualized and easily shared (25). To develop the *RLBase* TrackHub, the *buildGenomeBrowserHub.R* script from the *RLBase-data* GitHub repository was executed (see *Code availability*). Briefly, this script is used to build all the HTML descriptions for each human coverage track in *RLBase* along with a “oneFile” that ensures they will be accessed correctly in the Genome Browser session. The oneFile was also augmented to include the RL regions, along with regions of high G or C skew (see *Genome annotations*), and R-loop forming sequences (see *R-loop forming sequences*).

#### Running RLSeq and data upload

Finally, for each sample in *RLBase* which yielded called peaks, the *RLSeq* function was executed (see *RLSeq*) and HTML reports were generated. These steps were executed as part of running the *runRLSeq.R* script from the *RLBase-data* GitHub repository (see *Code availability*). The resulting *RLRanges* objects and *RLSeq* HTML reports were uploaded to the *RLBase* AWS S3 bucket along with all other processed data sets.

### RLPipes

RLPipes is a command-line interface (CLI) tool written in python and hosted on Bioconda (26) which is used for upstream processing of R-loop mapping data (see *Code availability*). It can accept FASTQ or BAM files and it can also be used with Sequence Read Archive (SRA) or Gene Expression Omnibus (GEO) accessions. The typical workflow for using *RLPipes* follows the *build, check*, *run* pattern.

#### Build

*RLPipes build* is a command which processes a “sample sheet” into a configuration file which can be used directly with the underlying workflow. When used with local files (FASTQ or BAM), the *build* command simply verifies that the files all exist, that all the supplied arguments and options are valid, and it finds the library type (single-end or paired-end) and read length for each sample (using the *pysam* package (27) for BAM files and the *pyfastx* package for FASTQ files (28)).

If the samples supplied are SRA or GEO accessions, then the *pysradb* python package (29) is implemented to query the SRA database and return the following:

1. The SRA experiment accession
2. The SRA study accession
3. The experiment title (author-submitted)
4. The organism taxonomy ID
5. The SRA run accession(s)
6. The library layout (i.e., paired-end or single-end sequencing)
7. The total number of bases in the run
8. The total number of spots in the run

The organism taxonomy ID is then used to query the available genomes list (see *Preliminary data preparation*) and return the most recent genome assembly for the organism to be added to the standardized catalog. Finally, the read length is calculated via floor division:

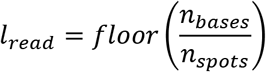

Where *l*_*read*_ is the read length, *n*_*bases*_ is the number of bases, and *n*_*spots*_ is the number of spots in the sequencing run. Finally, the configuration is saved for use with downstream functions.

#### Check

The *check* operation in *RLPipes* is a useful tool for verifying the configuration with the underlying *snakemake w*orkflow manager (30) (equivalent of using the *dry-run* and *dag* flags in *snakemake*). It also provides a visualization depicting all the jobs which will be run as part of the workflow.

#### Run

All steps in the processing pipeline are run automatically via the *snakemake* workflow manager (30) using pre-packaged *conda* environments. A typical analysis workflow will involve the following procedure: First, raw reads in SRA format are downloaded for each SRA run via the *prefetch* software from NCBI *sra-tools* (31). Then, reads are converted to “FASTQ” format using *fastq-dump* from *sra-tools* (31). Next, technical replicates are merged and interleaved (in the case of paired-end data) using *reformat.sh* from *bbtools* (32). Then, reads are trimmed and filtered with *fastp* (33), generating a quality report.

For R-loop mapping data, reads are aligned to the appropriate genome using *bwa mem* (34) or *bwa-mem2* (35), a faster implementation that users can enable with the *bwamem2* flag. Then, alignments are filtered (minimum quality score of 10 enforced), sorted, and indexed using *samtools* (36) and duplicates are marked using *samblaster* (37). Then, peaks are called using *macs3* (38) and coverage is calculated using *deepTools* (39). For RNA-Seq data, reads are aligned to the appropriate transcriptome using *salmon quant* (17), generating read count tables.

### RLHub

RLHub is an *ExperimentHub* (23) R/Bioconductor package which provides convenient access to the processed datasets available in *RLBase*. It is available in Bioconductor v3.14 and has built-in accessor functions for quickly downloading and caching data. To generate these datasets, the *prepRLHub.R* script from the *RLBase-data* GitHub repository was executed (see *Code availability*). Full documentation of all objects and accessor functions provided through *RLHub* is provided by the *RLHub* reference manual (see *Code Availability*).

### RLSeq

*RLSeq* is an R/Bioconductor package for downstream analysis of R-loop mapping samples. It is available via Bioconductor 3.14 and it depends upon the *RLHub* package. *RLSeq* contains three primary functions: (1) *RLRanges*, (2) *RLSeq*, (3) *report*. The following sections detail the functions in the *RLSeq* package.

#### RLRanges

The primary class structure used in *RLSeq* is the *RLRanges* object, initialized with the *RLRanges* function. This object is an extension of the *GRanges* class from the *GenomicRanges* R package (40) to provide additional validation and storage for the following slots:

1. Peaks (the *GRanges* containing R-loop mapping peaks)
2. Coverage (a URL or file path to a bigWig file)
3. Mode (the type of R-loop mapping)
4. Label (“POS” or “NEG”, the author-assigned label)
5. Genome (the UCSC genome ID)
6. Sample name (A sample name used for visualization and reporting)
7. *RLResults* (a list-like class for storing the results of *RLSeq* analysis)

#### Analyze R-loop forming sequences (*analyzeRLFS*)

R-loop forming sequences are regions of the genome with sequences that are favorable for R-loop formation (41). They are computationally predicted with the *QmRLFS-finder.py* software program (42) and serve as a test of whether a sample has mapped R-loops (11). The *analyzeRLFS* function provides a simple permutation testing method for analyzing the enrichment of RLFS within a provided peakset. The full analysis procedure is described in detail in our recent work (11).

#### Predicting sample condition (*predictCondition*)

Following R-loop forming sequences (RLFS) analysis, the quality model is implemented for predicting the sample condition (i.e., “POS” if the sample robustly mapped R-loops and “NEG” if the sample resembles a negative control). This is accomplished with the *predictCondition* function which performs all the steps described in detail our recent work (11) to render a prediction for each sample.

The results of this prediction, along with associated features and metadata, are stored in the *RLResults-predictRes* slot within the *RLRanges* object and returned to the user. For more detail, see the *RLSeq* reference manual (see *Code availability*).

#### Feature enrichment testing (*featEnrich*)

A custom list of R-loop relevant genomic annotations was curated for the human (hg38) and mouse (mm10) genomes (see *Genomic annotations*) and made available via the *RLHub* R/Bioconductor package (see *RLHub annotations*). In *RLSeq*, each annotation type is tested for enrichment within supplied *RLRanges*, yielding enrichment statistics. The procedure for this testing is described in detail our recent work (11).

#### Correlation analysis (*corrAnalyze*)

The *corrAnalyze* function performs a sample-level correlation test that can be used to assess sample-sample similarity by calculating coverage signal (from genomic alignments) around high-confidence R-loop sites (15). The *corrAnalyze* function ingests an *RLRanges* object (with a valid *coverage* slot) and performs the following procedure: (1) the coverage is quantified within the high-confidence sites and added as a column to the signal matrix (see *gs_signal* reference in the *RLHub* documentation to learn more about this matrix), and (2) then the *cor* function in R is used to calculate the Pearson correlation to yield a correlation matrix. The correlation matrix is saved in the *RLResults-correlationMat* slot of the *RLRanges* object and returned to the user.

#### Gene annotation (*geneAnnotation*)

The *geneAnnotation* function provides a simple procedure for annotating *RLRanges* peaks with gene IDs by overlap. Briefly, gene annotations are automatically downloaded using the *AnnotationHub* R package (43) and then overlapped with the ranges in the *RLRanges* object using *bed_intersect* from *valr* (21). The mapping between peak IDs and gene IDs is saved in the *RLResults-geneAnnoRes* slot of the *RLRanges* object.

#### R-loop regions overlap test (*rlRegionTest*)

R-loop regions (RL regions) are R-loop consensus sites generated during the *RLBase-data* workflow (see *R-loop Regions*). The *rlRegionTest* function uses a simple procedure for finding RL regions which overlap with peaks in the supplied *RLRanges* object. It also calculates the significance and odds ratio of the overlap using Fisher’s exact test implemented via the *bed_fisher* function from the *valr* R package (21). The results of this test are saved in the *RLResults-rlRegionRes* slot of the *RLRanges* object.

#### RLSeq

The primary workflow in the *RLSeq* package can be conveniently run in one step with the *RLSeq* function. This command will run, in order, (1) *analyzeRLFS,* (2) *predictCondition*, (3) *featureEnrich*, (4) *corrAnalyze*, (5) *geneAnnotation*, (6) *rlRegionTest*. The results are saved in the corresponding *RLResults* slots of the *RLRanges* object and returned to the user.

#### *RLSeq* plotting functions

*RLSeq* provides a variety of useful plotting functions which summarize and present analysis results:

1. *corrHeatmap* – generates a heatmap of the correlation matrix and annotations produced by *corrAnalyze* using either the *pheatmap* (44) or *ComplexHeatmap* (45) R packages.
2. *plotEnrichment* – generates hybrid violin-box plots showing the Fisher’s exact test odds ratio for each annotation calculated in *featureEnrich*. Importantly, it displays these results for the user-supplied sample alongside the distribution of results for the public samples within the *RLBase* database.
3. *plotRLRegionOverlap* – generates a Venn diagram showing the overlap between the user-supplied sample peaks and the R-loop regions, along with overlap statistics as calculated by the *rlRegionTest* function.
4. *plotRLFSRes* – plots the Z-score distribution or Fourier transform of the Z-score distribution in a metaplot, based on the analysis results from *analyzeRLFS*.

##### Report

The *RLSeq report* function ingests an *RLRanges* object which has already been processed by the analysis functions in *RLSeq*. It then uses RMarkdown templates to automatically build a user-friendly HTML report showcasing all results with summary tables and plots (see the *RLSeq* reference for an example HTML report).

### RLBase

*RLBase* is written in R shiny (46) and uses the *RLBase* Amazon Web Services (AWS) S3 bucket as a back-end storage solution. Notably, *RLBase* also includes extensive documentation which provide screenshots of all features with verbose descriptions, along with terminology explained, and FAQs. The following sections describe the features of *RLBase* and their methods.

#### RLBase datasets

*RLBase* contains standardized and reprocessed R-loop mapping data from 693 public samples (*RLBase* v1.0). The data were found via manual curation and reprocessed from raw sequencing reads using the *RLPipes* command-line tool (see *Code availability*). The data processing workflow is fully documented in our previous work (11). The resulting data was then uploaded to AWS S3 and made available publicly. The data and access methods are fully described in the “Downloads” section of the *RLBase* web server.

Following sample reprocessing, the data were quality-controlled using the quality control approach developed in our previous work (11) which classifies samples as “POS” (expected to map R-loops) or “NEG” (not expected to map R-loops). These classifications are also provided in the *RLBase* datastore and the models are accessible via the *RLBase* “Downloads” page and through the *RLHub* R package (see *Code availability*).

After quality control, high-confidence samples were analyzed to derive R-loop regions (RL regions), consensus sites of R-loop formation across the human genome. For details on the procedure used to produce these regions, see the description in our previous work (11). The RL regions can be downloaded via the *RLBase* “Downloads” page or through the *RLHub* R package (see *Code availability*).

#### Samples

The “Samples” page provides an interactive interface for exploring the 693 samples (RLBase v1.0) contained within *RLBase*. The interface for this tab is divided into three sections: (1) RLBase Samples Table, (2) Table Controls, (3) Outputs. The “RLBase Samples Table” is an interactive, searchable, sortable, paginated table built with the *datatables* JavaScript library (via the *DT* R package) (47). It contains metadata for every sample in RLBase. Selecting a row in the table will update the “Outputs” automatically. The “Table Controls” are interactive user-interface (UI) elements which control the data displayed in the table and in the outputs.

The coordination between UI elements, table row selection, and changes in the “Outputs” is due to the built-in reactivity of R *shiny* (46). Caching is also employed to improve the performance of this interface. Finally, the “Outputs” display a wealth of information, data, and plots that describe the samples (see *Results*, *RLBase*). The plots are produced with the built-in plotting functions in *RLSeq* (see *RLSeq plotting functions*) and with the *plotly* R package (48).

#### R-Loop Regions

The “R-Loop Regions” page provides an interactive interface for exploring the 64,418 R-loop regions (RL regions) uncovered in *RLBase* v1.0 (see *R-loop regions*). Like the “Samples” page, it includes three sections which interact via the *reactivity* within *shiny* (46): (1) RL Regions Table, (2) Table Controls, (3) Outputs. The “RL Regions Table” displays the RL regions and their metadata (see *R-loop Regions*). User-interface (UI) elements in the “Table Controls” filter the RL Regions Table based on criteria such as “Repetitive” (overlaps with repetitive elements). The “Outputs” show both an interactive summary of the RL region selected in the table, and a plot showing the relationship between RL region abundance (log2RLRPM) and RL region expression (log2TPM) (see *R-loop regions*). Finally, this page includes links to an interactive UCSC genome browser session which contains an interface for accessing all available coverage tracks and RL region signal tracks.

#### Analyze

The “Analyze” page provides an in-browser interface for analyzing R-loop mapping data. The API is simply the *RLSeq* R package but the interface in *RLBase* is designed to make these functions accessible to non-programmers. The sample entry form is used to describe the sample metadata and upload peaks in the manner necessary to construct an *RLRanges* base object (see *RLRanges*). Notably, no capability to upload coverage tracks is provided due to the size limitations of the server. Upon clicking the “Start” button, a background R process is launched using the *r_bg* function from the *callr* R package (49). Then, the following steps are performed: (1) A UUID (universally unique identifier, via the *uuid* R package (50)) is assigned. (2) The *knitr* R package (51) is used to render a template *Rhtml* which produces an HTML progress page that is uploaded to the AWS S3 location for the user’s sample and which the user is prompted to open. The progress indicator polls for updates using a custom *JavaScript* function and then it updates progressively as the analysis proceeds. (3) A sweetalert (via the *shinyWidgets* R package (52)) is displayed to alert the user that their sample is processing and to show them the link for viewing the progress page. (4) The *RLRanges* function from the *RLSeq* package (see *RLRanges*) is used to build the *RLRanges* object. (5) The *RLSeq* function is run to process all results for the sample (see *RLSeq*). (6) The *report* function is implemented to knit an HTML report (see *report*). (7) The finished *RLRanges*, *RLSeq* HTML report, and log files are all uploaded to the user’s directory (identified only by UUID) in the AWS S3 bucket. (8) The user is alerted via sweetalert that their sample is ready. If there was an error, *knitr* is used to knit the *Rhtml* error template and this is uploaded instead of the results page along with all log files to aid in debugging. For more information on the usage and results of this analysis, see the *Results-RLBase* section.

#### Download

The *RLBase* “Download” page provides convenient access to all processed data generated as part of the *RLBase-data* workflow (see *Code availability*). Instructions for bulk access via the AWS CLI are provided along with fine-grained access options for “Processed data files”, “RLHub downloads”, and “Raw and misc data”. The “Processed data files” tab provides an interactive, searchable table constructed with the *DT* R package (47) that provides download links for processed data files, such as peaks and coverage. The “RLHub downloads” panel provides a table constructed with the *kableExtra* R package (53) which includes direct links for downloading the data objects in *RLHub* as well as the functions which can be used to access them via the *RLHub* R package. The “Raw and misc data” panel provides an HTML-based guide to accessing raw and miscellaneous datasets.

#### Documentation

The documentation for *RLBase* was written in RMarkdown and knitted to an HTML file. The HTML is included in the *RLBase* web application via an *iframe*.

## Results

As described in our recent work (*Miller et al., 2021*), we processed and standardized 693 R-loop mapping samples via a purpose-built computational pipeline, *RLPipes* (**Fig. 1A**) (11). These data were further analyzed using *RLHub* and *RLSeq*, R/Bioconductor packages for downstream R-loop data analysis (**Fig. 1A**) (11). From meta-analysis of high-confidence R-loop samples, we derived R-loop regions (RL regions), sites of consensus R-loop formation (11). We now present *RLBase,* a user-friendly webserver which facilitates the exploration of public R-loop mapping samples and RL regions, access to standardized and reprocessed datasets, and the ability to analyze user-supplied R-loop mapping data in the browser (**Fig. 1B**). The following describes the primary features of *RLBase*, their usage, and the outputs which they generate.

**Figure 1.**
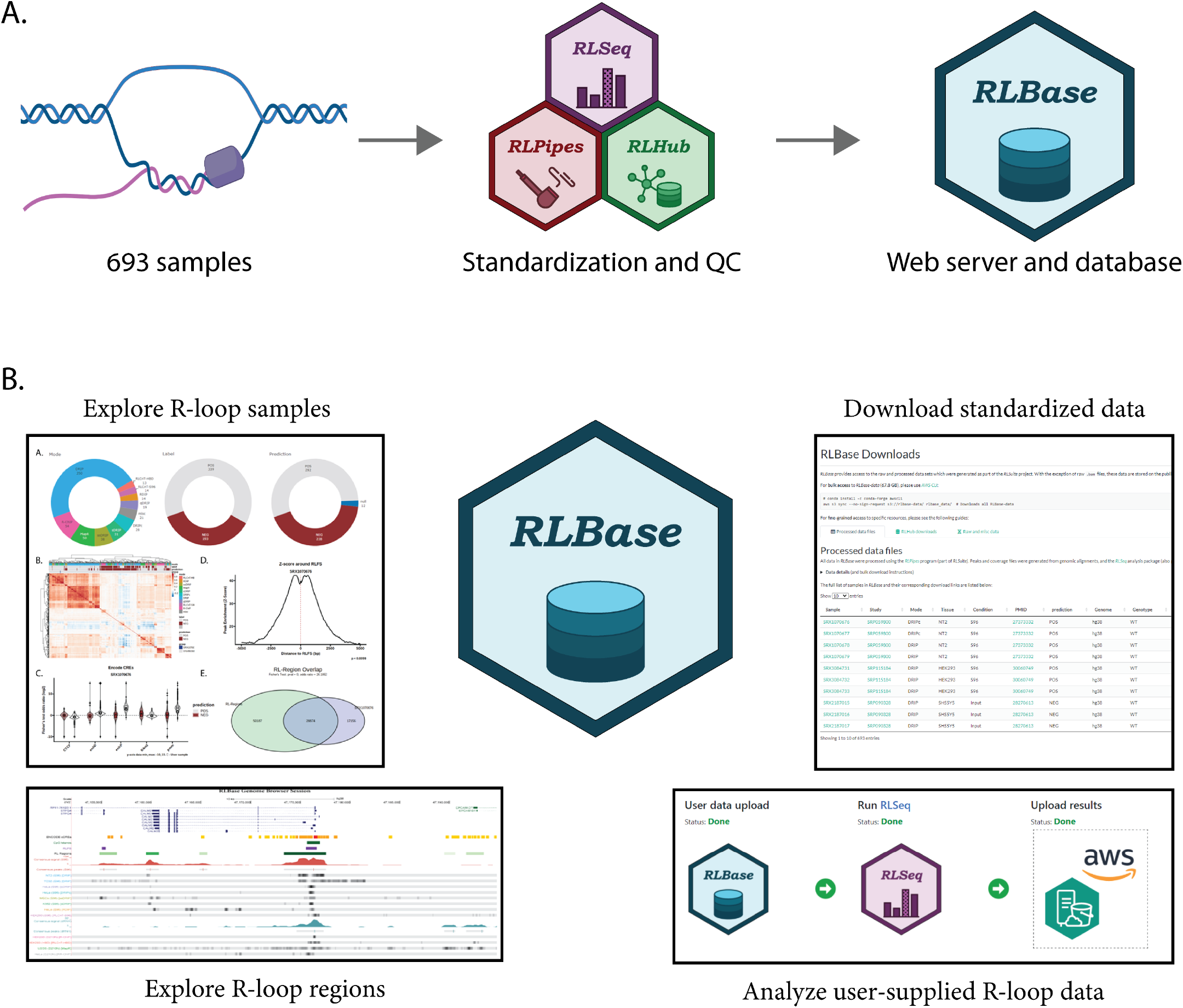
RLBase overview. (A) Graphical illustration showing the processing steps for RLBase. 693 R-loop datasets were downloaded and reprocessed using the RLPipes, RLHub, and RLSeq software as described in our recent work *(11)*. RLBase is a shiny web server which was developed based on these software and data. (B) Graphical illustration depicting the core functionality of RLBase. RLBase provides the capability to (1) explore R-loop mapping samples and generate summary visualizations, (2) explore R-loop regions and view them in the genome browser, (3) analyze user-supplied R-loop mapping data in the browser, and (4) download standardized and reprocessed R-loop mapping data.

### Samples

The “Samples” page (**Figure S1**) provides the capability to explore the 693 publicly available R-loop mapping datasets provided by *RLBase* (v1.0.0). As described in our recent work, these public R-loop mapping samples were reprocessed, standardized, quality controlled, analyzed with genomic feature enrichment analysis, and then used to derive R-loop regions (RL regions), sites of consensus R-loop formation (11). These analyses yielded a wealth of data which *RLBase* provides for exploration via the “Samples” page (**Figure S1**). Notably, this exploration is interactive as it contains user controls (**Fig. S1A**), an interactive data table (**Fig. S1B**), and responsive outputs (**Fig. S1C**) which provide the user with the capability to select samples and browse the visualizations and data relevant to them (**Figure 2**).

**Figure 2.**
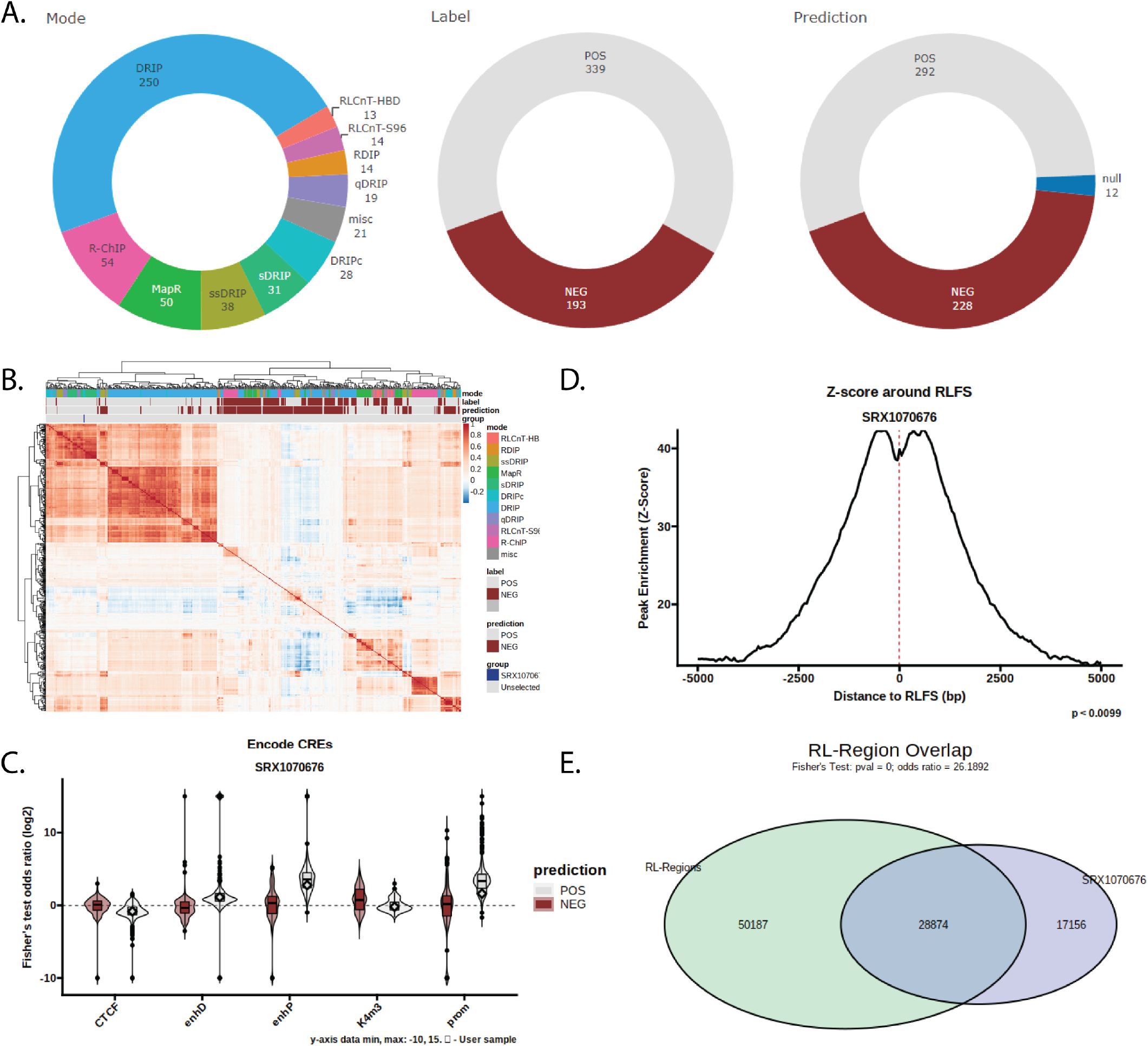
Select visualizations generated from RLBase. (A-E) Visualizations produced by RLBase with all modes, “Show labeled controls” and “Show predicted control”, and “hg38” genome options selected in the “Table controls”. Additionally, the DRIPc sample SRX1070676 was selected in the “RLBase Samples Table”. (A) Donut charts showing the proportions of samples by Mode, Label, and Prediction. (B) The sample-sample correlation heatmap from the “Sample-sample comparison” panel. (C) Genomic feature plots showing the distribution of ENCODE cis-regulatory element (CRE) feature enrichment within “POS” (predicted to map R-loops) and “NEG” (not predicted to map R-loops) samples. The user-selected sample (SRX1070676) is highlighted in each feature. Abbreviations: CTCF – CTCF binding site; enhD – distal enhancer; enhP – enhancer-promoter; K4m3 – H3K4me3 histone modification site; prom – promoter site. (D) R-loop forming sequences (RLFS) analysis plot with p value from permutation testing. (E) Venn diagram showing the overlap between selected sample ranges (SRX1070676 peaks) and RL regions. P value and odds ratio from Fisher’s exact test.

The sample page output panel (**Fig. S1C**) contains: (1) The “Summary” panel (**Figure S2**), (2) The “Sample-sample comparison” panel (**Figure S3**), (3) The “Annotation” panel (**Figure S4**), (4) The “RLFS” panel (**Figure S5**), (5) the “RL Regions” panel (**Figure S6**), and (6) the sample “Downloads” panel. The following will describe these outputs.

#### Summary

The summary panel provides a high-level overview of all samples in the “RLBase Samples Table” (**Fig. S2B-D**) and of the specific sample selected in that table (**Fig. S2A**). It is used for examining the representation of various R-loop mapping modes, labels, and predictions (**Fig. 2A**) among the samples selected via the “Table Controls” (**Fig. S1A**). From this interface, the user can also see the summary results for the sample which they select in the “RLBase samples table” (**Fig. S1A, S2A**). From these results, the user can easily explore the data at a high level and perform univariate analyses.

#### Sample-sample comparison

The next panel, “Sample-sample comparison” (**Figure S3**), provides the user with visualizations which reflect the similarities and differences between the samples selected in the “Table Controls” (**Fig. S1A**) and reveals the relationship between these samples and the single sample which is selected in the “RLBase Samples Table” (**Fig. S1B**). The primary visualizations are a sample-level heatmap (**Fig. 2B, S3A**) and a PCA plot (**Fig. S3B**). These plots use the Pearson correlation between R-loop mapping samples around high-confidence R-loop sites to reveal which samples are most similar with each other and which are most different. These visualizations can be used to examine how certain modalities differ from one another and showcase the “false positive” samples included in public R-loop mapping data (samples which are expected to map R-loops, but likely do not). Moreover, they allow users to explore how an individual sample of interest relates to all other samples within the selected data. This can be useful for evaluating how a particular sample relates to those it should be most alike.

#### Annotation

The “Annotation” panel (**Figure S4**) showcases the results obtained from running the *RLSeq featureEnrich* function for each sample in the dataset. The tabs in the output interface (**Fig. S4B**) control which annotations are displayed at any given time, the “Table Controls” (**Fig. S1A**) control the background data present in the plot, and the selected row in the “RLBase Samples Table” (**Fig. S1B**) controls which sample is highlighted in the plots (**Fig. S4C, 2C**). Finally, the “Split” selector controls whether the data is split by sample prediction, sample label, or whether there is no split (**Fig. S4A**). The plots display the distribution of feature enrichment results across all samples selected, expressed in terms of the log2 odds ratio from Fisher’s exact test (**Fig. 2C**). These plots allow the user to assess both the differences found within the dataset across annotations and a specific sample of interest which they want to observe annotations for. These results reveal novel relationships between R-loops and genomic features and provide a useful quality metric as we demonstrated in our recent work (11).

#### RLFS

R-loop forming sequences (RLFS) are genomic regions favorable to the formation of R-loops (16, 42). Moreover, they can be used to assess R-loop mapping sample quality, as we recently demonstrated in describing our RLFS analysis method and R-loop quality model (11). RLFS analysis was performed for each sample in *RLBase* via the *RLSeq analyzeRLFS* function. The resulting analysis plot, generated via the *RLSeq plotRLFSRes* function are provided for each sample in *RLBase* (**Fig. 2D, S5C**). The “RLFS” panel provides this plot along with other outputs which can be used to assess sample quality (**Figure S5**): (1) a permutation testing plot (**Fig. S5B**) as obtained from the *plot* method within the *regioneR* package, (2) an RLFS plot with Fourier transform applied (**Fig. S5D**), which pertains to the features used by the quality model to render a quality prediction, and (3) a summary of the results from quality analysis (**Fig. S5A**). These results together give the user the ability to assess the quality of each sample.

#### RL Regions

R-loop regions (RL regions) are consensus sites of R-loop formation discovered from the meta-analysis of high-confidence R-loop mapping samples, as we described in our recent work (11). The overlap of each sample in *RLBase* with these RL regions was calculated via the *RLSeq rlRegionTest* function. The visualization and full data table associated with these results is provided in the “RL Regions” panel (**Figure S6**). The visualization is a Venn diagram showing the overlap of the peaks in the user-selected sample with the RL regions (**Fig. 2E, S6A**). The p value and odds ratio are derived from the Fisher’s exact test. The RL region table is also provided (**Fig. S6B**), which lists all the RL regions uncovered by the peaks in the user-selected sample. These results provide the user with the capability to explore the degree of overlap between RL regions with the sample they select. It also allows the user to view the specific RL regions which were uncovered in the selected sample.

### R-Loop Regions

The “R-Loop Regions” tab (**Figure S7**) provides the tools necessary to explore consensus sites of R-loop formation (RL regions) derived from high-confidence *RLBase* samples, as described in our recent work (11). These regions contain a wealth of metadata, including their ID, genomic location, associated genes, and confidence level (**Fig S7B**). The user is provided with “Table Controls” (**Fig. S7A**) that allow them to control which RL regions are displayed and how gene names are shown. The “All genes” checkbox allows the user to control whether all genes overlapping an RL region are displayed, or whether only well-annotated genes are shown (those which appear in pathway databases). The “Repetitive regions” checkbox controls whether to show RL regions which overlap with repetitive genomic sequences, such as pericentromeric regions. Finally, the “Correlated with expression” checkbox will, if selected, only show the RL regions for which a significant correlation between R-loop abundance and gene expression was observed. The output panel (**Fig. S7D**) displays a summary of the RL region currently selected in the “RL Regions Table” (**Fig. S7B**) and displays a correlation plot showing the relationship between R-loop abundance and gene expression levels.

Finally, the *RLBase* UCSC genome browser session is provided (**Figure 3, S7C**). This browser session provides access to all the human R-loop mapping samples in *RLBase* along with consensus signal, peaks, RL regions, RLFS, and other relevant annotations. Taken together, these features provide the user with the capability to explore RL regions, observe their association with expression, and view them alongside all other datasets in *RLBase* in the UCSC genome browser.

**Figure 3.**
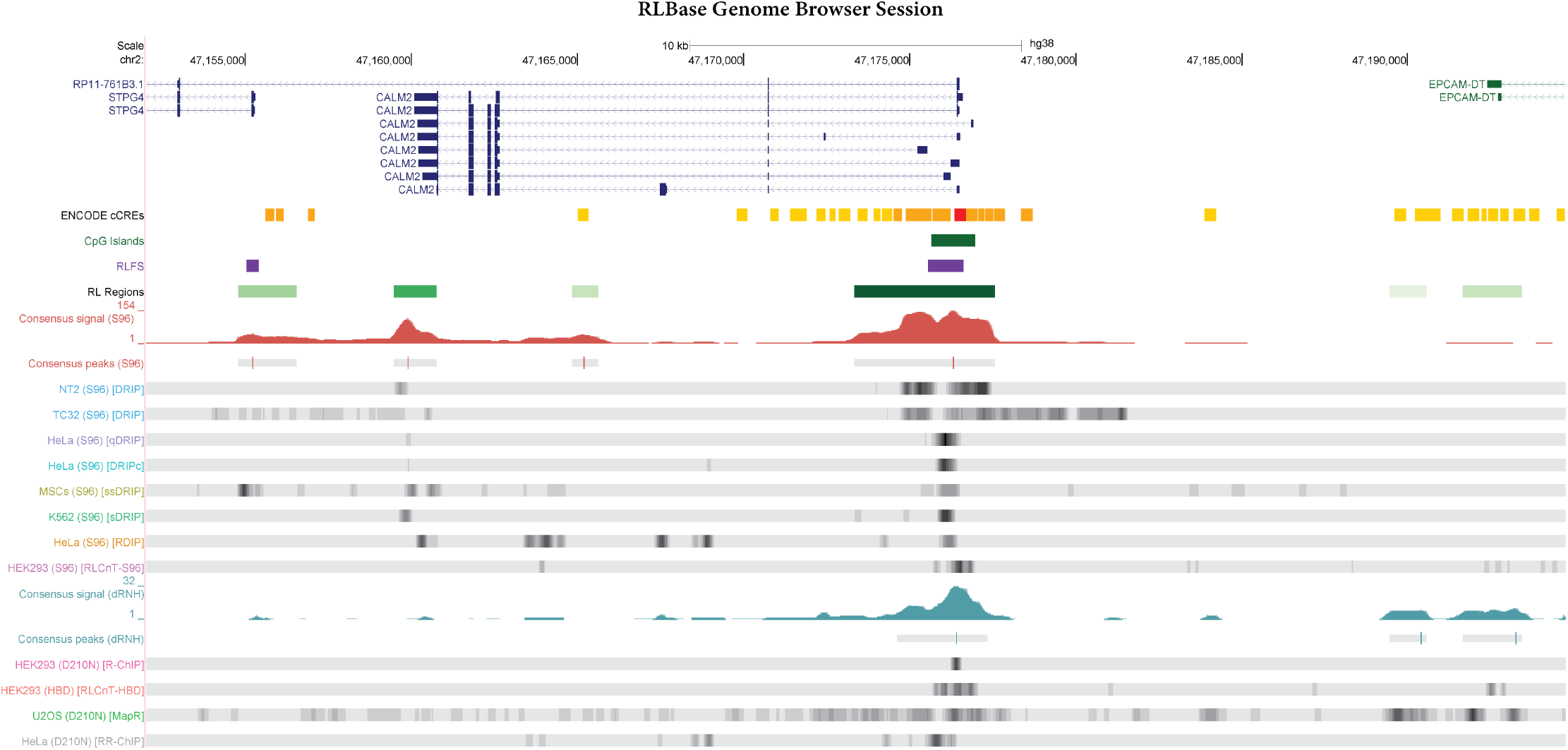
RLBase genome browser session. The browser session includes representative RLBase sample coverage tracks alongside S9.6 and dRNH consensus signal, RL regions, CpG islands, R-loop forming sequences (RLFS), and ENCODE cis regulatory elements (CREs). The screen capture shown here was taken in the area surrounding the CALM2 gene.

#### Analyze

The “Analyze” page (**Figure S8**) provides users with the ability to analyze their own R-loop mapping data in the browser without the need for bioinformatics skill. To analyze their data, a user provides metadata about the sample to be analyzed (**Fig. S8A**), the peaks (in BED format) for the sample (**Fig. S8B**), and agrees to the privacy policy (**Fig. S8C**). Having performed these steps, the user selects “Start” and launches the analysis job. The user is presented with the results link which will show the progress of the analysis and, when completed, will update to show the analysis results (**Figure S9**). The results page contains a verbose description of the analysis (**Fig. S9A**), the progress indicator (**Fig. S9B**), and a table with sample metadata and the results (once available) (**Fig. S9C**). These results contain the *RLSeq* report which details the analysis results, the *RLRanges* R data object which the user can load in an R session for further analysis, and the logs generated by the analysis, which are useful for debugging should an error be encountered. Taken together, the “Analyze” page provides users with an easy and convenient in-browser interface to the *RLSeq* functions used for analysis, yielding quality reports which have a permalink suitable for sharing.

#### Download and Documentation

The “Download” page provides direct links to download all processed files associated with *RLBase*, including verbose descriptions of each item and of programmatic methods for accessing these data. It also includes instructions for how to download the alignment (BAM) files for each sample and how to access processed datasets via the *RLHub* R/Bioconductor package. Finally, the “Documentation” page provides verbose descriptions of features and addresses terminology and technical concepts which may need additional explanation.

## Conclusion

*RLBase* is a user-friendly web server for the exploration and analysis of R-loop data. It extends accessibility to the novel methods and data resources we developed as part of our recent work (11) by providing the capability to (1) explore 693 reprocessed and standardized R-loop mapping datasets, (2) explore R-loop regions (consensus sites of R-loop formation), (3) analyze user-supplied R-loop datasets, and (4) download all the processed and standardized data generated as part of our previous work (11). Moreover, the *RLSuite* software collection (*RLPipes, RLHub, RLSeq,* and *RLBase*) provides a robust infrastructure for R-loop bioinformatics.

## Code Availability

All software developed as part of this project are available publicly under MIT license. Purpose-built software packages used include (1) RLSeq (available via Bioconductor) - https://bioconductor.org/packages/devel/bioc/html/RLSeq.html, (2) RLHub (available via Bioconductor) - https://bioconductor.org/packages/devel/data/experiment/html/RLHub.html, (3) RLPipes (available via Bioconda) - https://anaconda.org/bioconda/rlpipes, (4) RLBase (available via GitHub) - https://github.com/Bishop-Laboratory/RLBase.

Additionally, the scripts used for preprocessing of all data provided in *RLBase* are available from the *RLBase-data* GitHub repository: https://github.com/Bishop-Laboratory/RLBase-data.

## Data Availability

All data are made available through the *RLHub* R/Bioconductor package and the *RLBase* web interface. *RLBase* URL: https://gccri.bishop-lab.uthscsa.edu/rlbase/. *RLHub* URL: https://bioconductor.org/packages/devel/data/experiment/html/RLHub.html.

## Acknowledgements

We want to thank the Bishop Laboratory for their insightful comments and feedback on the web server interface. We want to thank Lori Kern at Bioconductor for her review of *RLSeq* and *RLHub*. We want to also thank Simon Bray at Bioconda for his review of *RLPipes*. Figure 1A was created, in part, using BioRender.com.

