## Supplemental Figures for "Exploration and analysis of R-loop mapping data with *RLBase*"

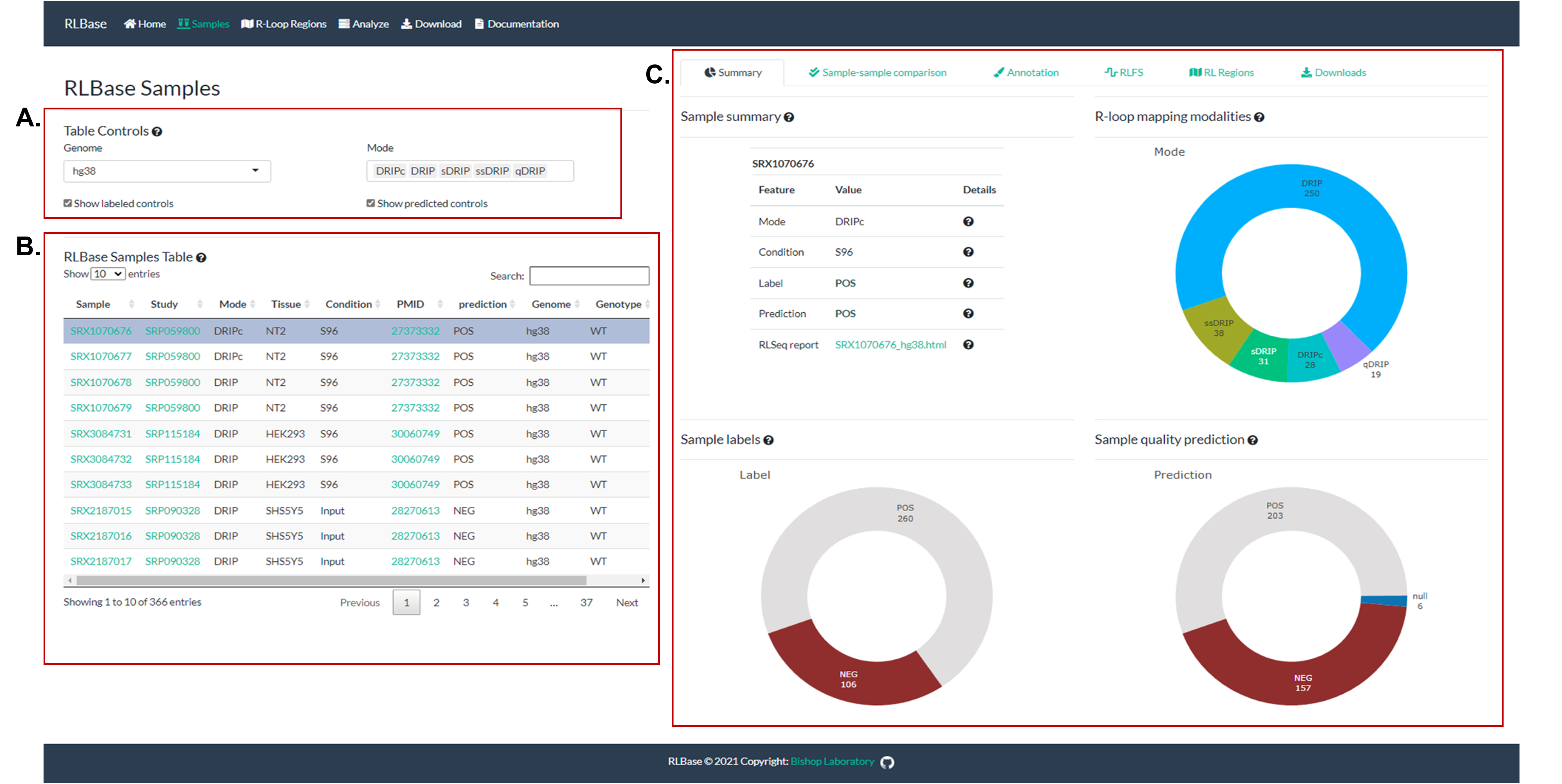

**Figure S1. The RLBase "Samples" page.** (A) Controls for the ‘RLBase Samples Table’ which modify the displayed data in (B) and (C). (B) The ‘RLBase Samples Table’ which displays data about the R-loop mapping samples in RLBase. Selecting an individual row will change the output in (C). (C) Output pane showing the data and plots relevant to the selected sample(s) in the table.

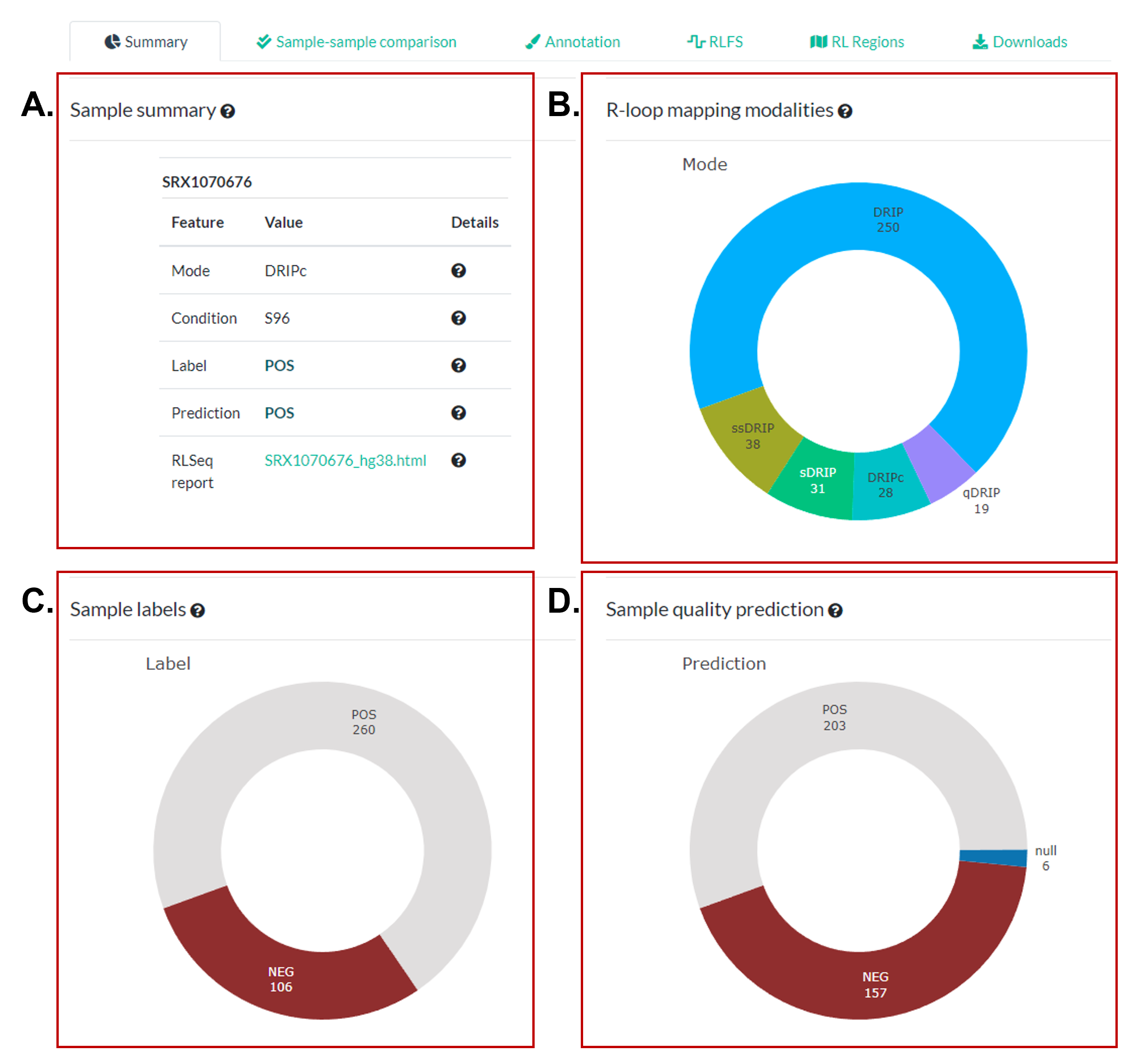

**Figure S2. The “Summary” panel.** (A) Summary characteristics of the selected sample. (B) The representation of R-loop mapping modalities within the selected data sets (determined by the “Table Controls”). (C) Similar to (B) showing the representation of sample labels (“POS” meaning expected to map R-loops, “NEG” meaning not expected to map R-loops). (D) Similar to (C) except that “POS” and “NEG” are predicted by the ML quality model (See model details).

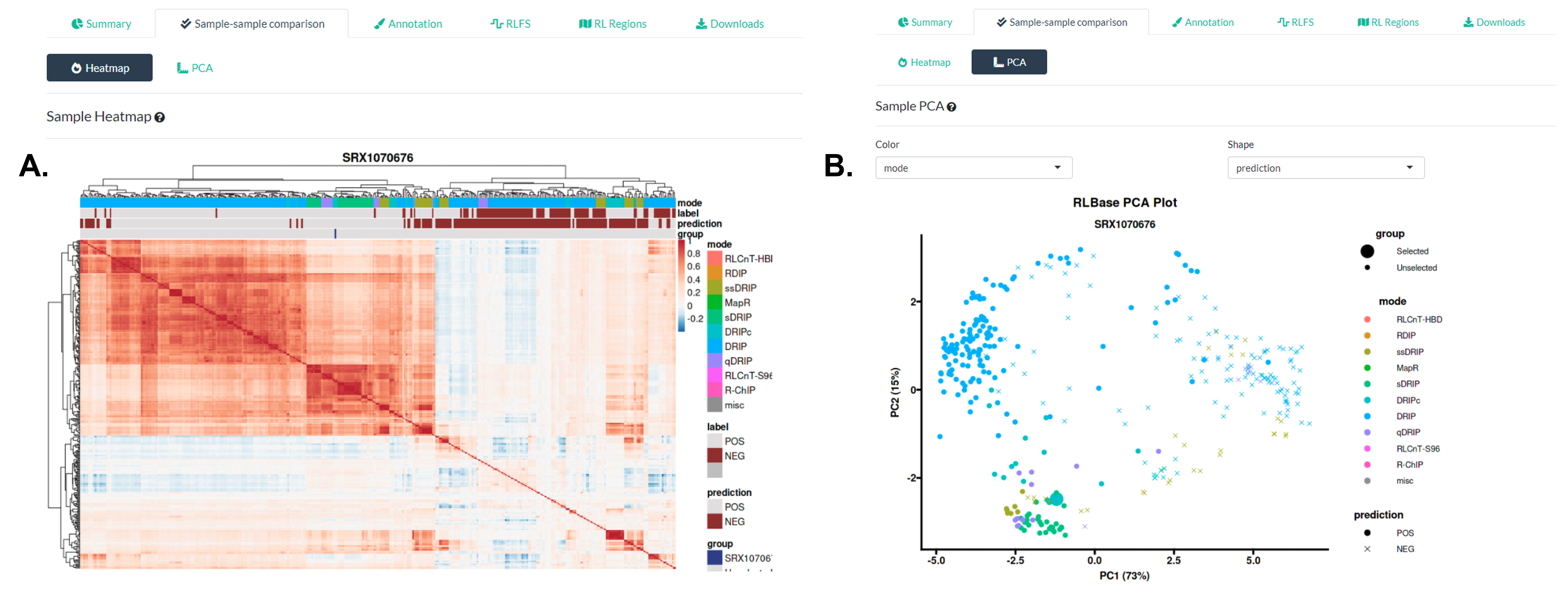

**Figure S3. The “Sample-sample comparison” panel.** (A) A heatmap showing sample-sample pearson correlation of coverage (bigWig) tracks around gold-standard R-loops (See here for additional detail). (B) Similar to (A) but with a scatter plot showing PC1 and PC2 from principal component analysis (PCA) of the correlation matrix. Controls allow for customization of the plot.

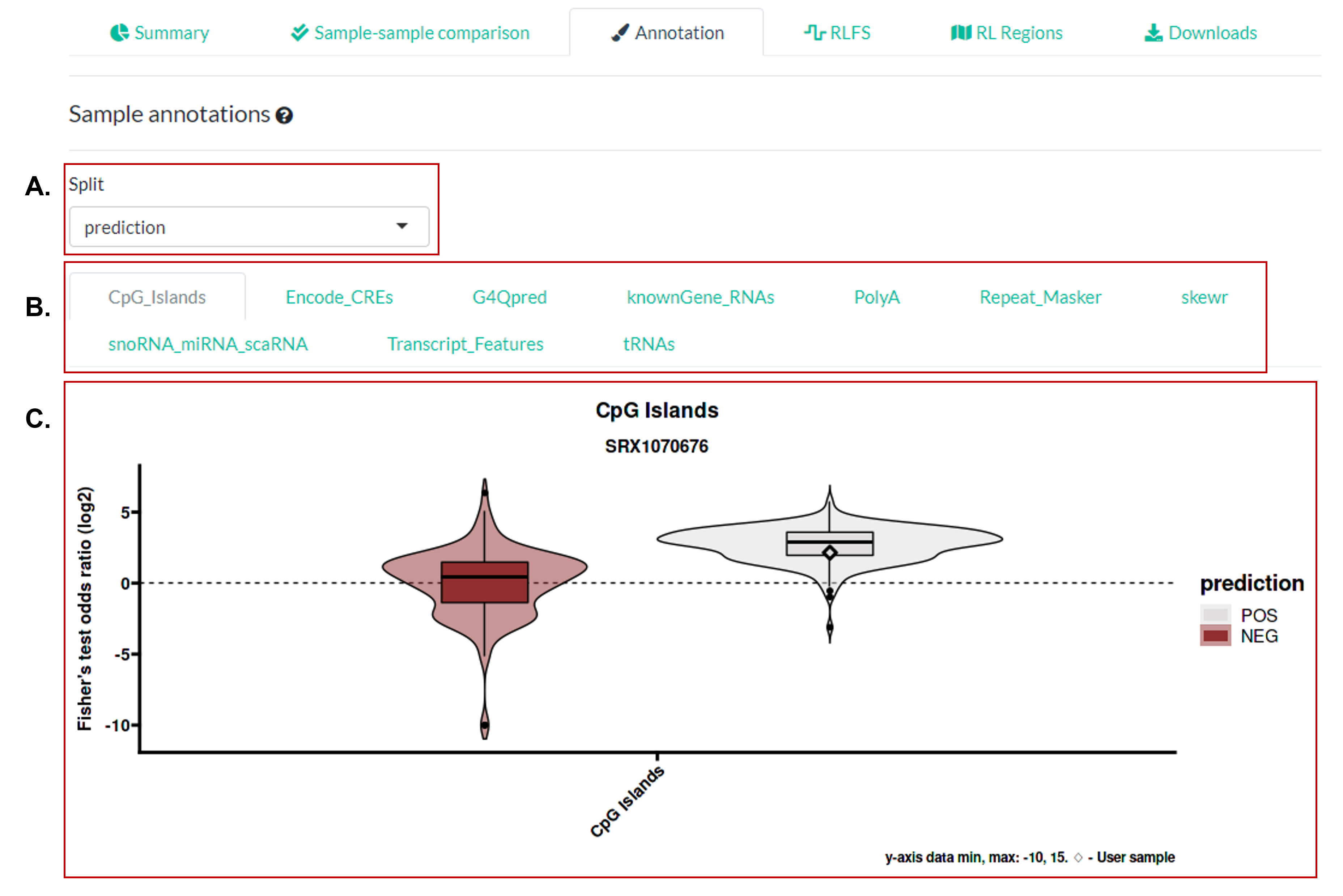

**Figure S4. The “Annotation” panel.** (A) A “Split” control that determines plotting in (C). (B) The databases for which plots are available. See the documentation of annotation databases here for more detail. (C) A plot showing the distribution of enrichment results (represented by the log2 Fisher’s exact test odds ratio). The sample selected in the table is shown as a diamond (if results are available). Additional information can be found in the RLSeq Vignette

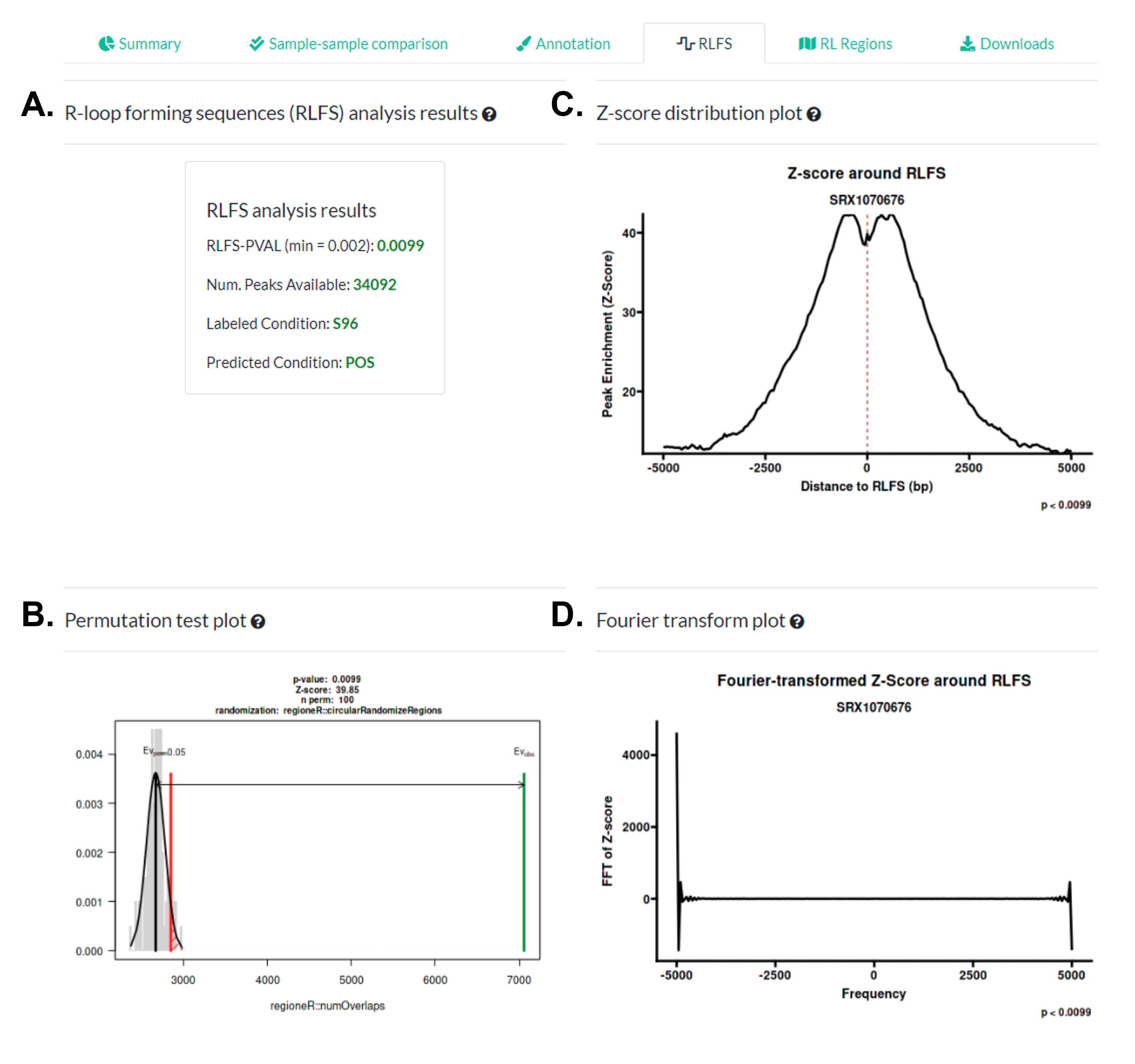

**Figure S5. The “RLFS” panel.** Additional details regarding the R-loop forming sequences (RLFS) analysis approach are described in the RLSeq vignette. (A) A summary of the results from the R-loop forming sequences (RLFS) analysis. (B) The empirical distribution plot from permutation testing. The distribution represents the overlap of RLFS with randomized ranges, whereas the green line represents the actual number of overlaps. (C) The Z-score distribution plot shows the Z-score of the non-random overlaps compared to random ranges within 5kb upstream and downstream from RLFS. This distribution is what the ML model uses to predict quality (along with the P Value). (D) The Fourier transform of (C).

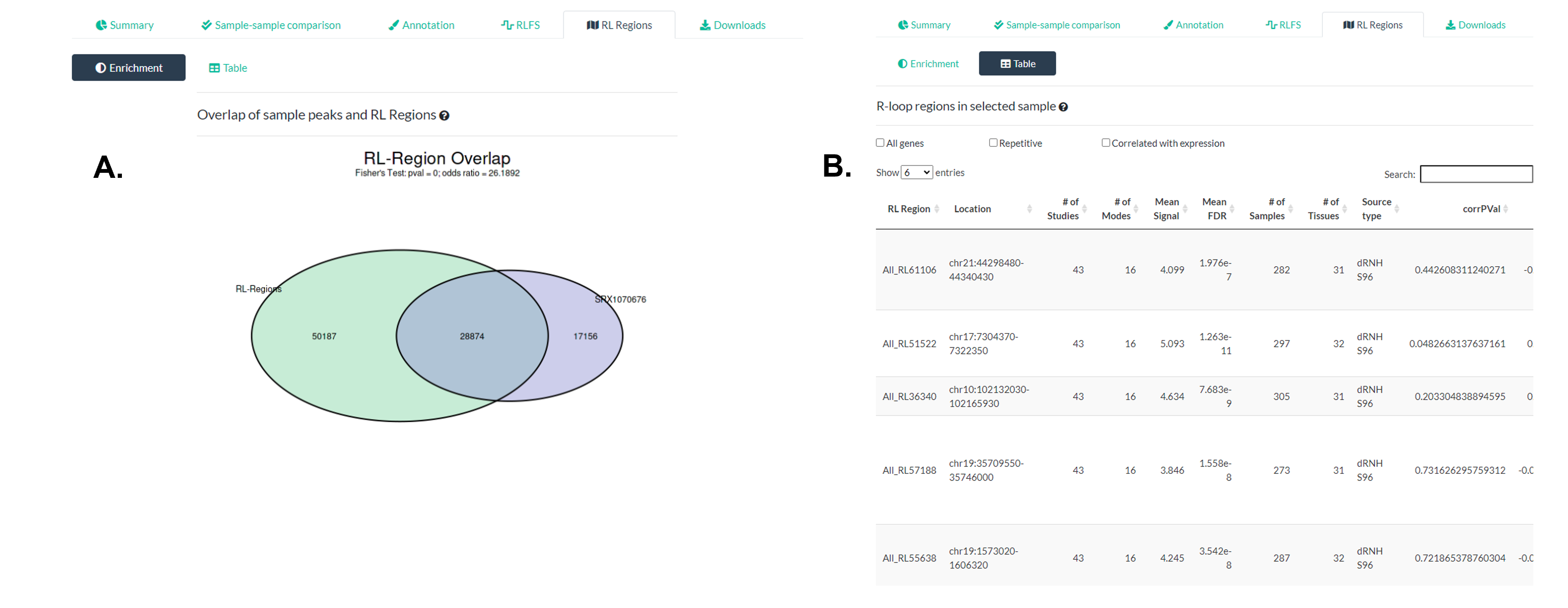

**Figure S6. The “RL Regions” panel.** R-loop regions (“RL Regions”; consensus sites of R-loop formation) are described in greater detail in the RLHub documentation. (A) The overlap of RL Regions and the peaks within the selected samples. The p value and odds ratio from Fisher’s exact test are also shown. See also the relevant section of the RLSeq vignette. (B) The table of RL Regions overlapping with the select sample’s peaks. See the RL Regions section for a detailed explanation of this interface.

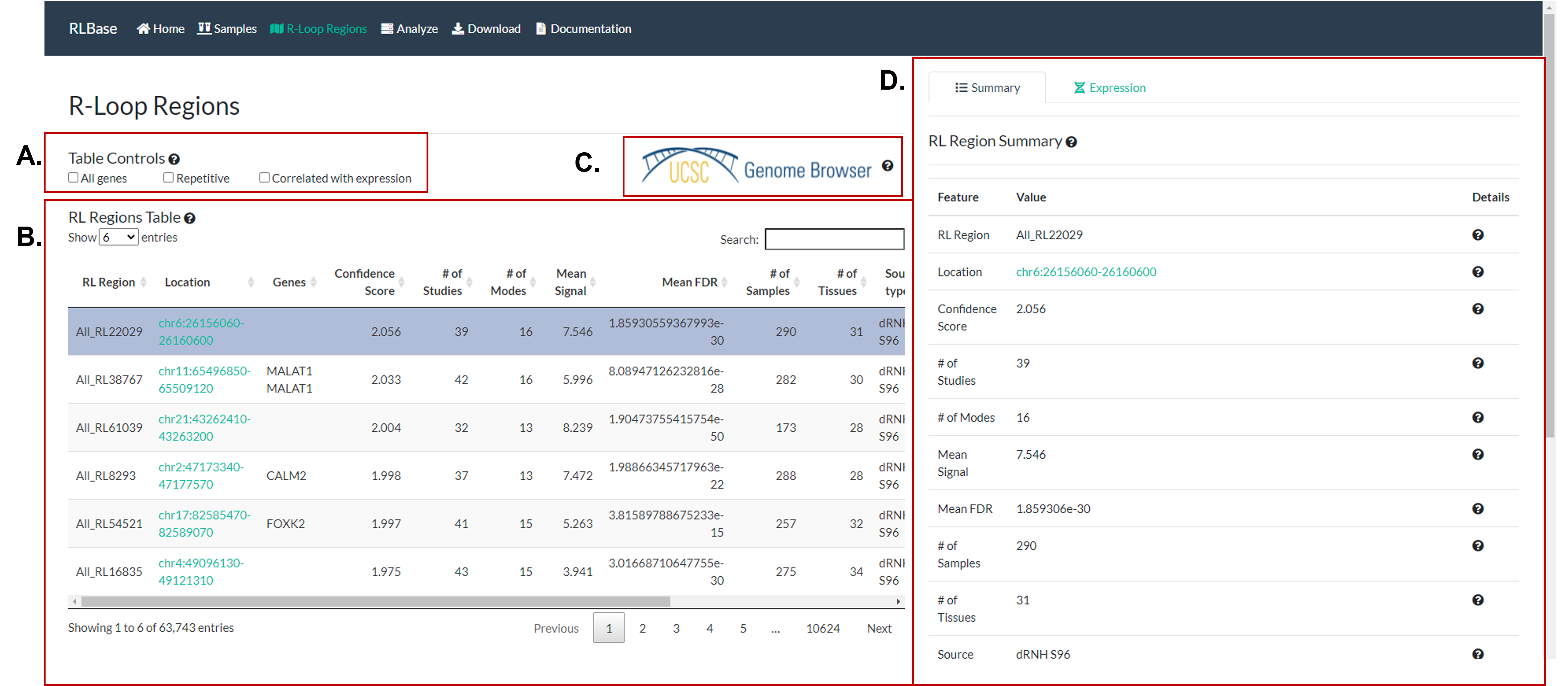

**Figure S7. The “R-loop Regions” page.** R-loop regions (“RL Regions”; consensus sites of R-loop formation) are described in greater detail in the RLHub documentation. This page provides an interface for exploring these data. (A) Controls for altering the data shown in (B). “All genes,” if selected, forces (B) to also show pseudogenes, RNA genes, and predicted genes. If “Repetitive” is selected, RL Regions that overlap with repetitive elements will be displayed. If “Correlated with expression” is selected, only the RL Regions with significant correlation between R-loop signal and expression will be shown. (B) The table of RL Regions along with metadata. The RL Regions in this table are controlled by (A) and selecting rows will impact the output in (D). For a full description of columns, please see the relevant RLHub documentation. (C) A link to the UCSC genome browser session for RLBase. This session contains all the coverage files for every human R-loop mapping sample, along with the consensus R-loop regions. (D) The output panel impacted by the selection in (B). See (Figure 8) for a full description.

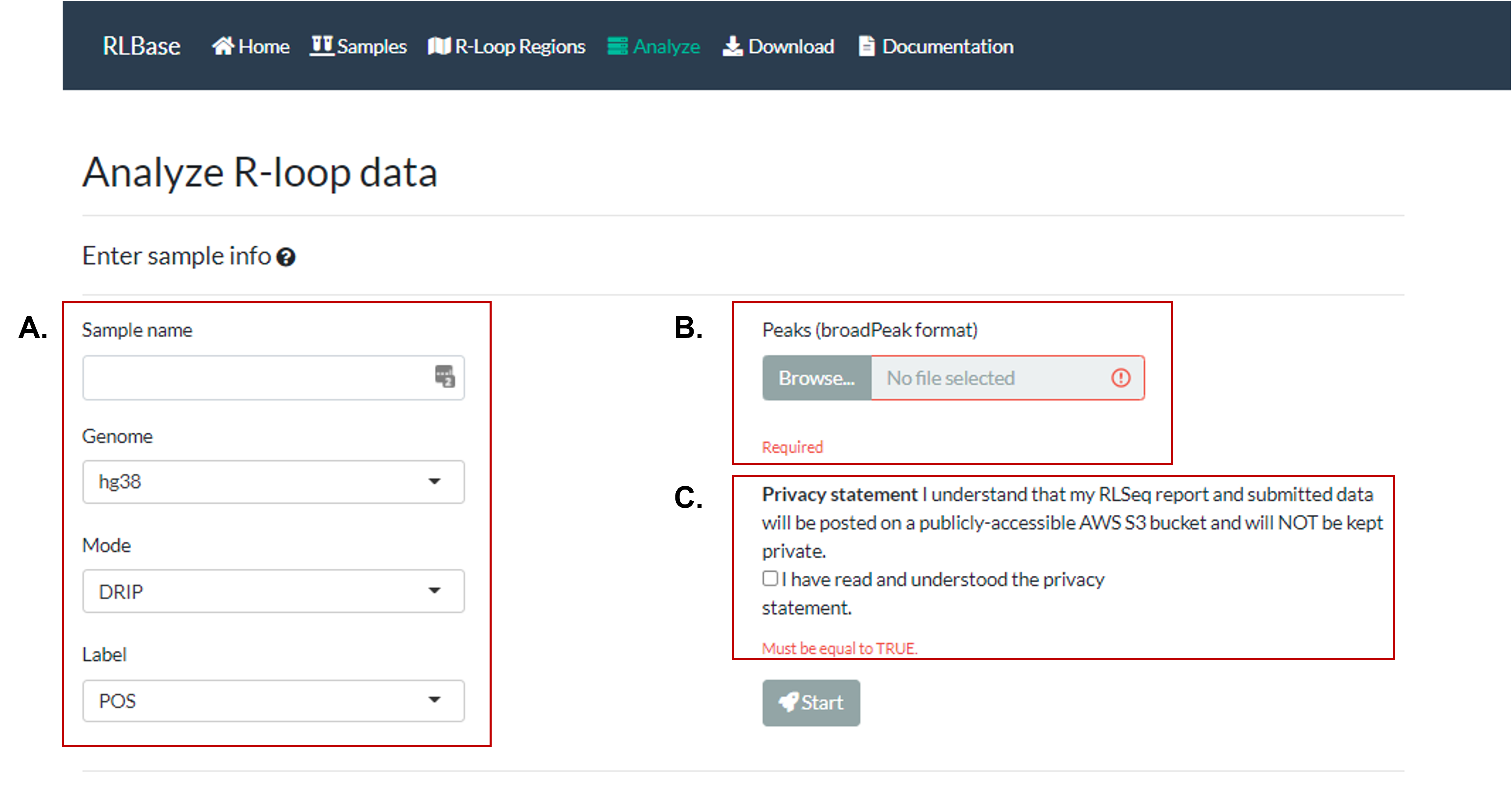

**Figure S8. The “Analyze” page allows users to analyze an R-loop mapping sample in the browser.** (A) Inputs describing the R-loop mapping sample. It is recommended to lift-over your data to “hg38” or “mm10” if possible, as RLSeq is most compatible with these genomes. For an explanation of the “Mode” and “Label,” please see the terminology section. (B) An upload input for the peaks to analyze (should be in broadPeak format if possible (narrowPeak and BED can also be used, but may produce unexpected results). (C) The user must also acknowledge the privacy statement, which specifies that uploaded data will be made publicly available in the analysis results link.

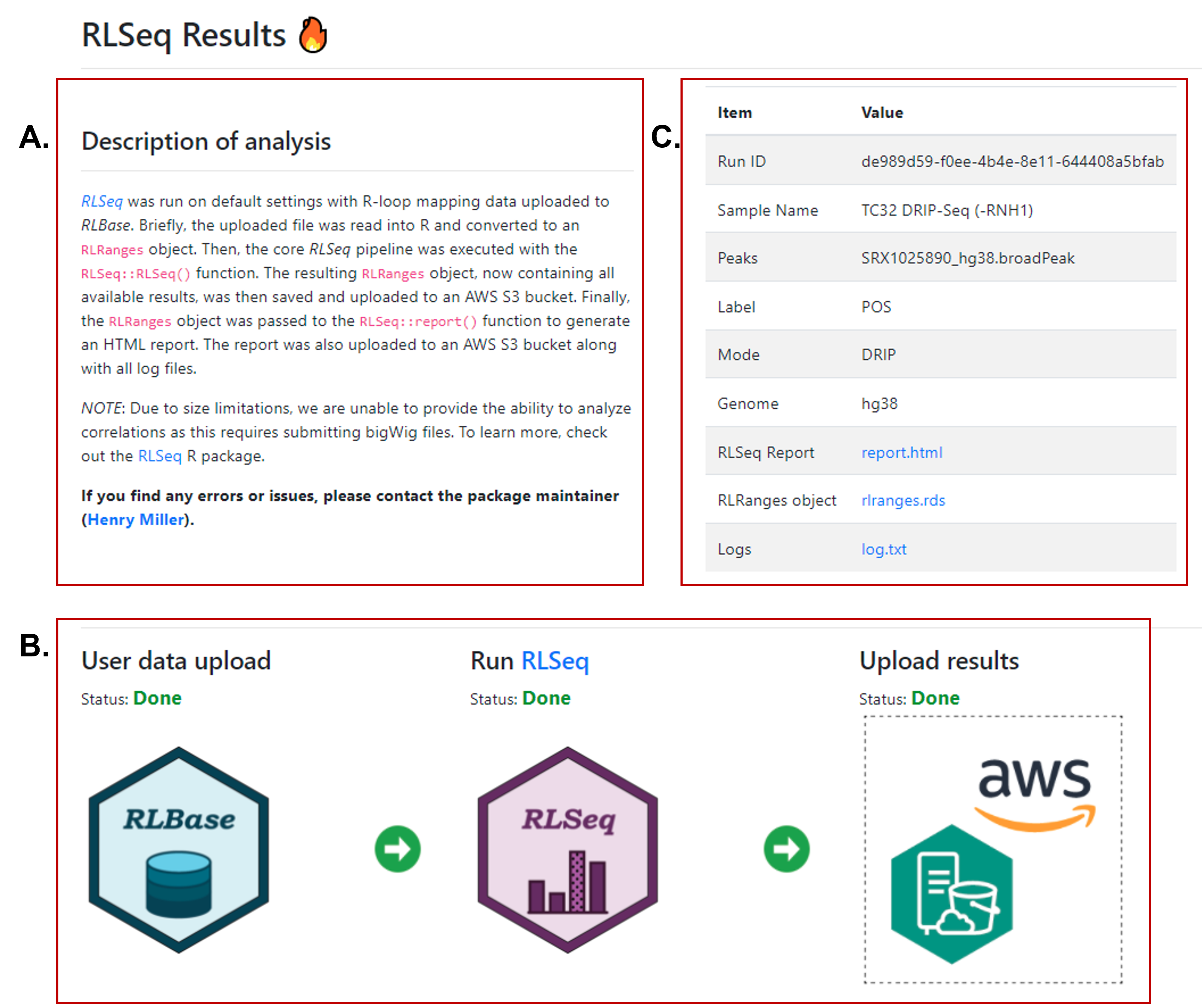

**Figure S9. The “RLSeq Results” page displays the analysis description and results (it also displays progress during the analysis).** (A) A verbose description of the analysis, recommended for citing RLSeq. See the RLSeq vignette for additional detail. (B) The progress indicators that display the current stage of the analysis. (C) Metadata about the uploaded sample and links to the RLSeq report, RLRanges R data object for the sample, and the log files.
